# MitoMAMMAL: a genome scale model of mammalian mitochondria predicts cardiac and BAT metabolism

**DOI:** 10.1101/2024.07.26.605281

**Authors:** Stephen P. Chapman, Theo Brunet, Arnaud Mourier, Bianca H. Habermann

## Abstract

Mitochondria perform several essential functions in order to maintain cellular homeostasis and mitochondrial metabolism is inherently flexible to allow correct function in a wide range of tissues. Dysregulated mitochondrial metabolism can therefore affect different tissues in different ways which presents experimental challenges in understanding the pathology of mitochondrial diseases. System-level metabolic modelling is therefore useful in gaining in-depth insights into tissue-specific mitochondrial metabolism, yet despite the mouse being a common model organism used in research, there is currently no mouse specific mitochondrial metabolic model available. In this work, building upon the similarity between human and mouse mitochondrial metabolism, we have created mitoMammal, a genome-scale metabolic model that contains human and mouse specific gene-product reaction rules. MitoMammal is therefore able to model mouse and human mitochondrial metabolism. To demonstrate this feature, using an adapted E-Flux2 algorithm, we first integrated proteomic data extracted from mitochondria of isolated mouse cardiomyocytes and mouse brown adipocyte tissue. We then integrated transcriptomic data from *in vitro* differentiated human brown adipose cells and modelled the context specific metabolism using flux balance analysis. In all three simulations, mitoMammal made mostly accurate, and some novel predictions relating to energy metabolism in the context of cardiomyocytes and brown adipocytes. This demonstrates its usefulness in research relating to cardiac disease and diabetes in both mouse and human contexts.

## Introduction

Mitochondria are essential organelles found in almost all eukaryotic cells and are indispensable for cellular bioenergetics, metabolism and homeostasis. One of their main objectives is to produce ATP through oxidative phosphorylation (OXPHOS). OXPHOS occurs within the inner mitochondrial membrane, where electrons are shuttled along an Electron Transport Chain (ETC) mediated by the mobile electron carriers, Coenzyme Q (CoQ) and cytochrome C. Electron transfer through each complex is coupled to proton translocation from the mitochondrial matrix to the intermembrane space which generates a Proton Motive Force (PMF) across the inner membrane that is used by ATP-synthase, to phosphorylate ADP to ATP (Hatefi, 1985). Several other metabolites are also directly oxidised by the ETC by reducing CoQ. In mammals, these include the mitochondrial glycerol-3-phosphate dehydrogenase (G3PDH)(Mráček *et al*., 2013), dihydroorotate dehydrogenase (DHODH)(Molinié *et al*., 2022), proline (Tanner *et al*., 2018; Pallag *et al*., 2022) and the electron transfer flavoprotein dehydrogenase (Lenaz *et al*., 2007), the first step of mitochondrial fatty acid oxidation.

Apart from the production of cellular energy, mitochondria are integral to various cellular and metabolic processes including pacing organism-specific development rates (Diaz-Cuadros *et al*., 2023), apoptosis (Suen *et al*., 2008), calcium signalling (Nicholls, 2005) and regulating ROS production, which itself is an important secondary messenger (Forman *et al*., 2010). Mitochondria also generate metabolic intermediates crucial for biosynthetic pathways and redox regulation (Borst, 2020). Due to their central roles in cellular metabolism, signalling and bioenergetics, dysregulated mitochondrial metabolism is associated with various human diseases, emphasising their critical role in maintaining cellular health (Liesa *et al*., 2009). Understanding the intricacies of mitochondrial metabolism is therefore essential for advancing knowledge of cell biology, physiology and medicine.

### Tissue-specificity of mitochondrial structure and content

Mitochondrial structure (Kuznetsov *et al*., 2009; Liesa *et al*., 2009), and proteome content vary across tissues (Hansen *et al*., 2024; Calvo and Mootha, 2010; Williams *et al*., 2018). Considering the metabolic roles played by proteins, proteomic changes would reroute metabolism to sustain different biological objectives in various cellular contexts. Therefore, mitochondrial metabolism and function are highly specialised to meet diverse cellular functions and bioenergetic needs. This is strongly evidenced in cardiomyocytes which are responsible for the control of the rhythmic beating of the heart and rely heavily on ATP to achieve maximal cardiac output (Karbassi *et al*., 2020). Brown Adipose Tissue (BAT), is a specialised type of adipose tissue with unique mitochondrial properties that permit thermogenic heat generation (Flatmark and Pedersen, 1975; Takeda *et al*., 2023). One key characteristic of brown adipocyte mitochondria is a high abundance of uncoupling protein 1 (UCP1), which is responsible for uncoupling OXPHOS from ATP production (Wang *et al*., 2019; Hansen *et al*., 2024). This uncoupling leads to the dissipation of the PMF across the inner mitochondrial membrane as heat, a process crucial for non-shivering thermogenesis (Nicholls, 1979, 2021), thus highlighting an alternate biological objective of the mitochondria within BAT. A better understanding of mitochondrial metabolism could, for instance, help reduce the prevalence of metabolic diseases in cardiac and other chronic metabolic diseases like diabetes.

### System-level modelling to gain in-depth insight into tissue-specific mitochondrial metabolism

Systems-level modelling of mitochondrial metabolism is essential to provide novel and testable model-driven insights into mitochondrial function and disease (Ben Guebila and Thiele, 2021; Heinken *et al*., 2021; Wagner *et al*., 2021; Tomi-Andrino *et al*., 2022). Flux Balance Analysis (FBA) is a computational method that implements linear programming in conjunction with a metabolic reconstruction to predict metabolic fluxes on the systems level (Orth *et al*., 2010; Sahu *et al*., 2021; Westerhoff, 2023). By integrating existing knowledge of mitochondrial biology into such a modelling framework, researchers can specifically analyse mitochondrial metabolism. Omics data, such as transcriptomics or proteomics can be integrated into a metabolic reconstruction using a variety of methods such as E-Flux2 (Kim and Lun, 2014; Kim *et al*., 2016), to produce context-specific metabolic models. Thus, metabolic modelling can facilitate a better understanding of the metabolic differences between tissues or disease conditions.

### The mitoCore model of human mitochondrial metabolism of cardiomyocytes

The mitochondrial metabolism of humans and mice is included in several metabolic reconstructions. Recon 1 was the first generic human metabolic model (Duarte *et al*., 2007) and from this, an orthologous mouse model (iMM1415) was then created (Sigurdsson *et al*., 2010). Recon 1 has been updated to Recon R2, which included additional biological information and the correction of various modelling errors such as Recon 1’s inability to correctly predict realistic ATP yields (Thiele *et al*., 2013). Recon R2 was subsequently upgraded to Recon 3D, which includes a total of 13,543 metabolic reactions and extensive human GPR associations (Brunk *et al*., 2018). From this model, an updated mouse model, iMM1865 was produced using a top-down orthology-based methodology by mapping human genes of Recon 3D to mouse genes (Khodaee *et al*., 2020).

One challenge facing predictive modelling at the genome-scale level however is that large models encompassing thousands of reactions are more prone to error than smaller and more concise models. This is a consequence of missing knowledge and annotation. Sources of error include incorrect parameterisation of reaction directionality constraints and issues relating to the incorrect compartmentalisation of reactions and metabolites. Using large genome-scale models to specifically predict mitochondrial metabolism can therefore result in mispredictions (Bi *et al*., 2022; Fritzemeier *et al*., 2017; Karp *et al*., 2023; Ong *et al*., 2020).

Several concise models of human mitochondrial metabolism exist (Smith and Robinson, 2011; Smith *et al*., 2017), with mitoCore representing the latest and most comprehensive model of human cardiomyocyte mitochondrial metabolism (Smith *et al*., 2017). MitoCore includes the ETC within its reconstruction and can accurately model the PMF associated with ATP production. This model has successfully been applied to model fumarase deficiency (Smith *et al*., 2017), impaired citrate import (Majd *et al*., 2018) and predicted accurate respiratory quotients on glucose and palmitate substrates (Cabbia *et al*., 2021) which demonstrates mitoCore’s potential to model human cardiomyocyte mitochondrial metabolism.

Mice are often employed as a model organism in mitochondrial research due to their highly similar structure, function and genetic homology with human mitochondria. This similarity makes mice a valuable model system for advancing our understanding of mitochondrial biology, mitochondrial dysfunction and disease, and for exploring potential interventions for mitochondrial-related disorders in humans. Because of these similarities, mitochondrial metabolism is routinely compared to human metabolism in diverse biological contexts (Porter *et al*., 2016; Diaz-Cuadros *et al*., 2023). Despite the prevalence of mice *in vivo*, *in vitro* and *in silico* models, there are no concise *in silico* models of mouse mitochondrial metabolism.

To address this limitation, and to valorise the opportunity presented by mitochondrial similarity, in this work, we have created mitoMammal, a mitochondrial metabolic network which can be used for constraint-based metabolic modelling of human and mouse mitochondria. Importantly, mitoMammal can be contextualised with -omics data emerging from humans or mice, allowing for the capacity to model the metabolism of both species. To demonstrate this novelty, we have integrated mitochondrial transcriptomic data from Brown Adipocytes (BAs), and then mitochondrial proteomic data from mice BAT and cardiac tissues. We found that integrating proteomic and transcriptomic data from humans and mice into mitoMammal predicted proline dehydrogenase and G3PDH reduction of CoQ, the export of hexadecanoic acid from BAT tissue, and glycine import to sustain cardiomyocyte metabolism.

## Methods

### Conversion of mitoCore into mitoMammal

To build the mitoMammal mitochondrial metabolic model, we identified the mouse orthologs of mitoCore’s gene-product-reaction (GPRs) using BioMart (Kasprzyk, 2011) and the ENSEMBL database (Martin *et al*., 2023), as well as orthology information stored within MitoXplorer2 (Marchiano *et al*., 2022). This resulted in 389 mouse orthologs out of the original complement *of* 391 MitoCore genes) (Supplementary Table S1). The set of mitoCore GPR rules was compiled to their corresponding logical expressions for mitoMammal based on orthology relations between human and mouse genes. A summary of mitoMammal construction is represented in Figure 1a (see also Supplementary Table S1). Gene modifications in the mouse version of the metabolic model reconstruction are listed in Table 1.

**Figure 1:**
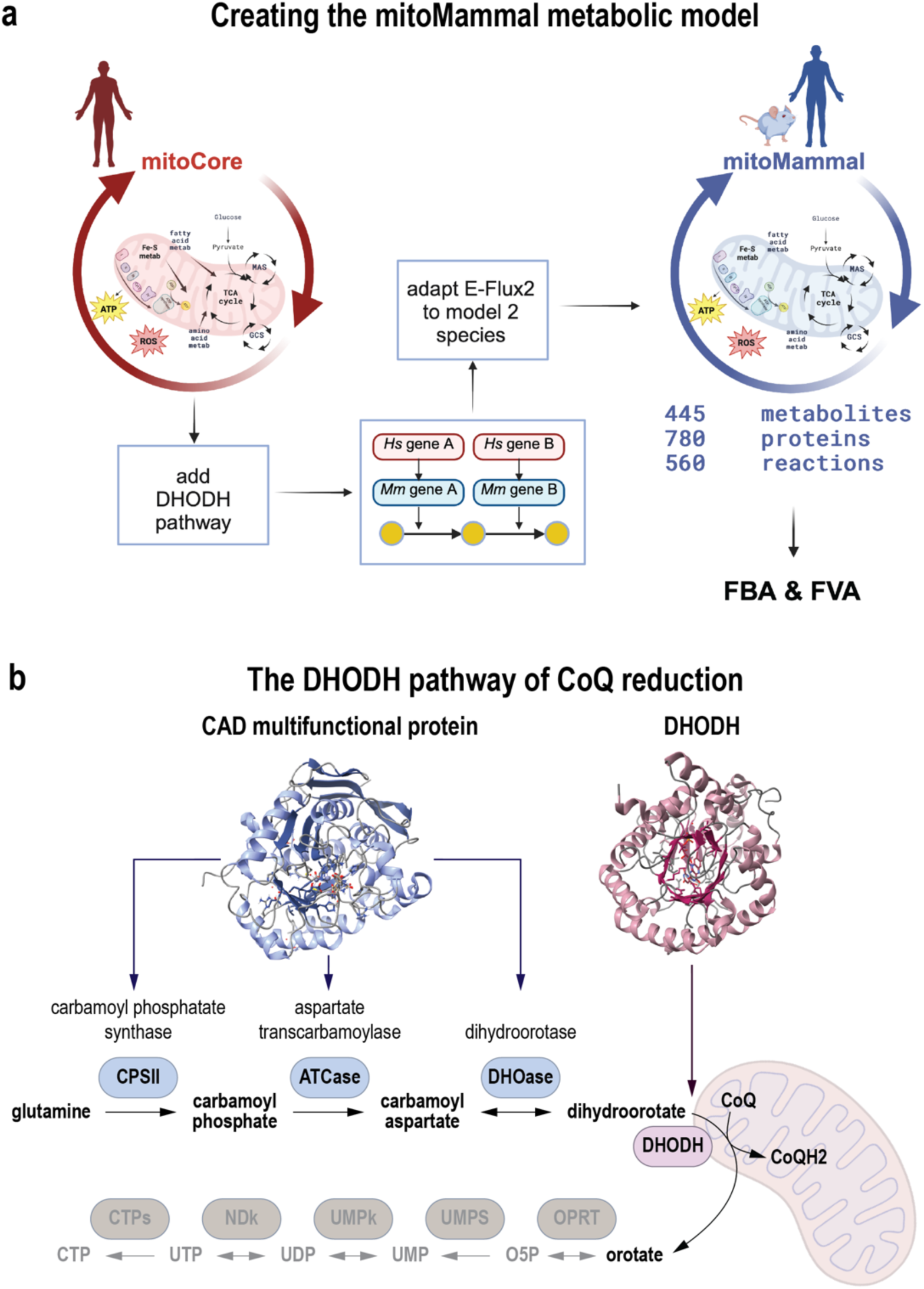
**(a**) Workflow for the construction of the mitoMammal metabolic network. **(b)** The DHODH pathway of CoQ reduction was added to the mitoMammal mitochondrial metabolic model.

**Table 1:**
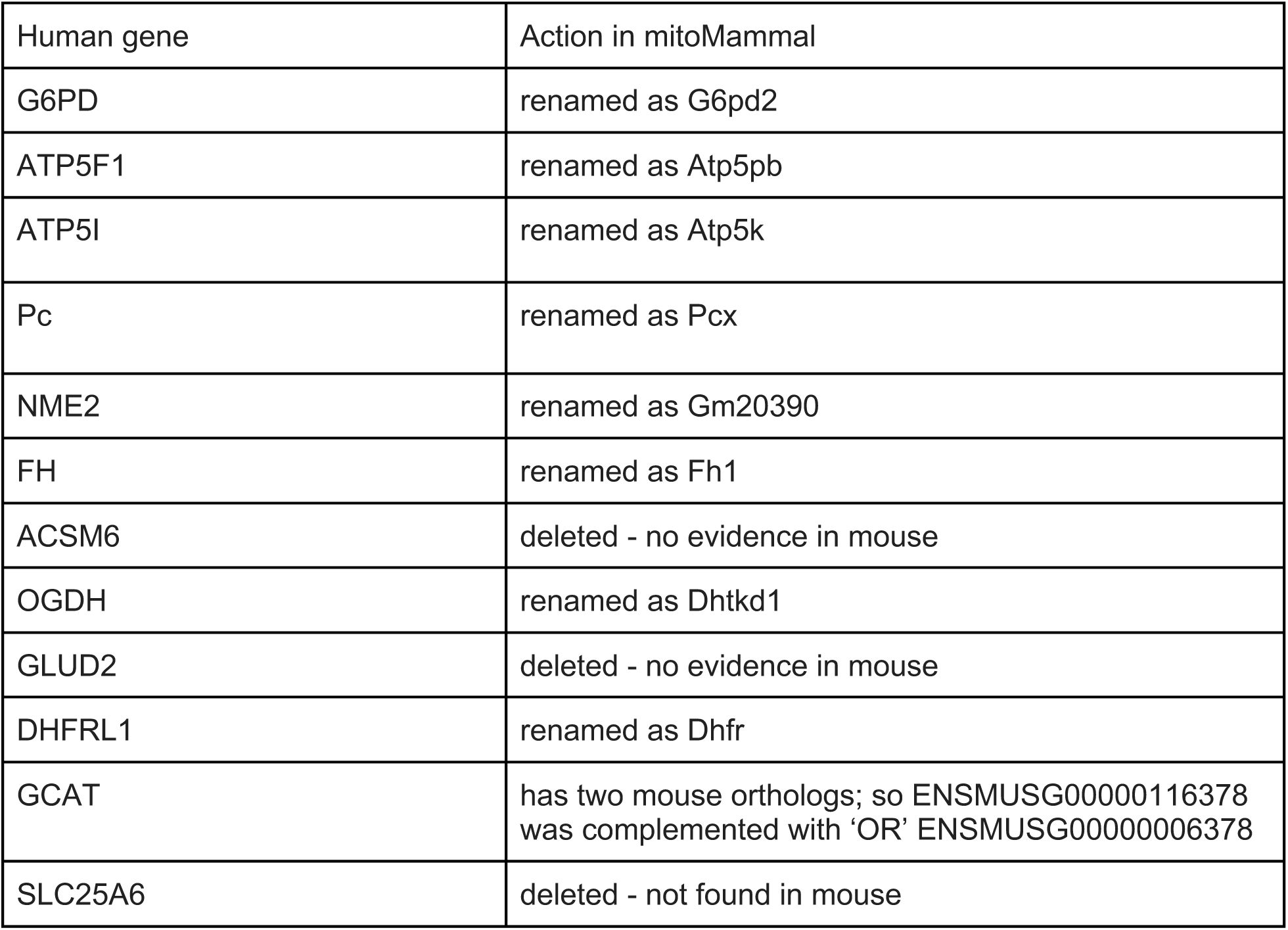
Changes applied from human to mouse in mitoMammal.

### DHODH expansion

The discovery that dihydroorotate can reduce CoQ in mouse mitochondria suggests that this is a conserved feature of all mammalian mitochondria (Molinié *et al*., 2022). MitoCore (Smith *et al*., 2017) was missing the reduction of CoQ by DHODH within the *de novo* pyrimidine synthesis pathway, while it contained glutamine metabolism, which is the starting substrate for this pathway. Initially, glutamine is converted to carbamoyl phosphate facilitated by carbamoyl phosphate synthase. Carbamoyl phosphate is then metabolised to carbamoyl aspartate through the activity of aspartate carbamoyltransferase, which is subsequently metabolised into dihydroorotate by the enzyme dihydroorotase (Figure 1b). In mammals, these three enzymes are part of a single multifunctional protein abbreviated as CAD (Carbamoyl Aspartate Dihydroorotase). Dihydroorotate then reduces CoQ to produce orotate, facilitated by the enzyme dihydroorotate acid dehydrogenase (DHODH) that sits at the surface of the outer mitochondrial membrane. As such, orotate is never imported into the mitochondria and remains cytoplasmic (Zhou *et al*., 2021). We included these metabolic reactions and new metabolites in mitoMammal. Orotate removal from the model was implemented by the addition of a demand reaction to maintain flux consistency. In total, five new reactions were added that incorporate four new metabolites and two new genes.

### Update to SBML3 (version 1)

The original MitoCore model was encoded using SBML level 2 annotation. We updated the mitoMammal to the most recent, relevant specification of SBML level 3 (version 1) (Keating *et al*., 2020) and validated the model for correctness using the online SBML validation tool (https://synonym.caltech.edu/validator_servlet/index.jsp; (Lister *et al*., 2007)).

### Adapting the E-Flux2 algorithm for mitoMammal

Because the MitoMammal metabolic model can integrate -omics data from two, instead of one species, we modified the E-Flux2 algorithm by allowing the user to select the organism the data originates from. The adapted algorithm then uses -omics data to constrain the reactions specific to the chosen species. All other features of the original E-Flux2 method were maintained as in the original description of the algorithm (Kim *et al*., 2016). The adapted E-Flux2 algorithm was used to constrain mitoMammal with mouse proteomic data and transcriptomic data from humans.

#### Mouse proteomic data

For mouse simulations, we integrated proteomic data from a recent study that extracted the mito-proteomes of isolated mitochondria from a range of mouse tissues (Hansen *et al*., 2024). Normalised protein counts of cardiac and brown adipose tissue (BAT) were scaled between 0 and 1. For cardiac tissue, we optimised ATP hydrolysis and for BAT simulations, we optimised the UCP1 reaction.

#### Human transcriptomic data

We used normalised RNA-sequencing data from Rao (Rao *et al*., 2023)(GEO dataset GSE185623) to model an *in vitro* differentiated hiPSC-derived BA. Normalized read counts were scaled between 0 and 1. The UCP1 was chosen to be optimised considering the essential role that UCP1 plays in uncoupling ETC from ATP synthesis in BAs.

### Flux Balance Analysis

Parsimonious FBA was performed using Python (version 3.8.5) in conjunction with the COBRApy toolbox (Ebrahim *et al*., 2013), using the default ‘GLPK’ solver.

The mitoMammal metabolic model, along with Jupyter notebooks and data used in this work are available at: https://gitlab.com/habermann_lab/mitomammal.

## Results

### The mitoMammal metabolic network for human and mouse mitochondrial metabolism

This work aimed to produce a generic mammalian metabolic model of mitochondrial metabolism that incorporates new knowledge on CoQ fueling. We first translated the genes from the human mitoCore model into mouse genes using orthology inference to create the basic mitoMammal model. Key metabolic pathways that include the TCA cycle, the Malate Aspartate Shuttle (MAS); OXPHOS and ATP synthesis; the Glycine Cleavage System (GCS), the proline cycle and fatty acid oxidation were also retained from the original model. MitoMammal now includes *de novo* pyrimidine synthesis from glutamate leading to the reduction of the CoQ complex by the enzyme DHODH. MitoMammal contains 780 genes encoding 560 metabolic reactions that involve 445 metabolites. The complete lists of reactions, metabolites, and associated fluxes from each simulation are available in Supplementary Table S1a, Supplementary Table S1b and Supplementary Table S1c respectively. The core metabolism and bioenergetics with associated import/export reactions of the model are depicted in Figure 2.

**Figure 2:**
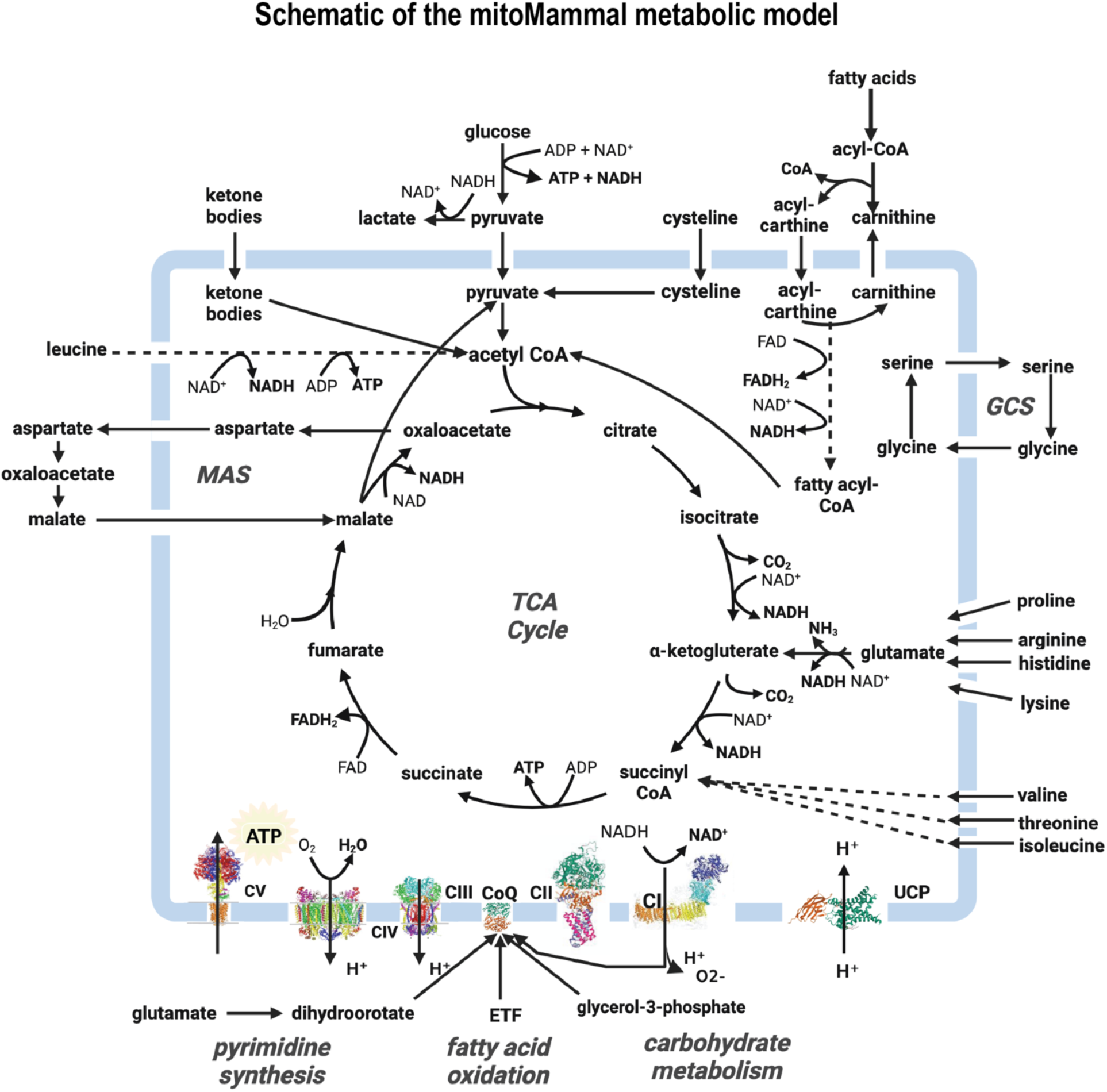
MitoMammal metabolic reconstruction consists of 780 genes (human and mouse orthologs) encoding 560 metabolic reactions. Initially constructed from mitoCore (Smith *et al*., 2017), it was expanded to include DHODH reduction of COQ and then supplemented with mouse orthologous genes. MitoMammal can be used for detailed mitochondrial metabolic studies of both human and mouse.

MitoMammal is based on MitoCore, a human specific cardiomyocyte mitochondrial model. MitoMammal was first tested on its ability to correctly produce accurate ATP levels from glucose oxidation. All nutrient input reactions except glucose and oxygen were constrained to zero to reflect aerobic glycolytic conditions. Maximisation of ATP hydrolysis was used as the objective function for these simulations and the model was then optimised using parsimonious FBA for all simulations reported in this work. As expected, MitoMammal correctly predicted the production of 31 molecules of ATP from 1 molecule of glucose (Supplementary Figure S1).

### Modelling cardiac mitochondrial metabolism by integrating mouse proteomic data of cardiac tissue

To demonstrate mitoMammals ability of modelling mouse cardiac mitochondrial metabolism, we integrated proteomic data harvested from mitochondria isolated from mouse cardiac tissue and optimised ATP hydrolysis. This resulted in 330 constrained reactions out of the complement of 560 reactions. In satisfying the objective subject to these constraints, MitoMammal predicted the import of ɑKG, H2O, glucose, oxygen, malate, oxaloacetate and glutamine, and the export of citrate, isocitrate, lactate, hydrogen, CO2, NH4, alanine and succinate (Figure 3a).

**Figure 3:**
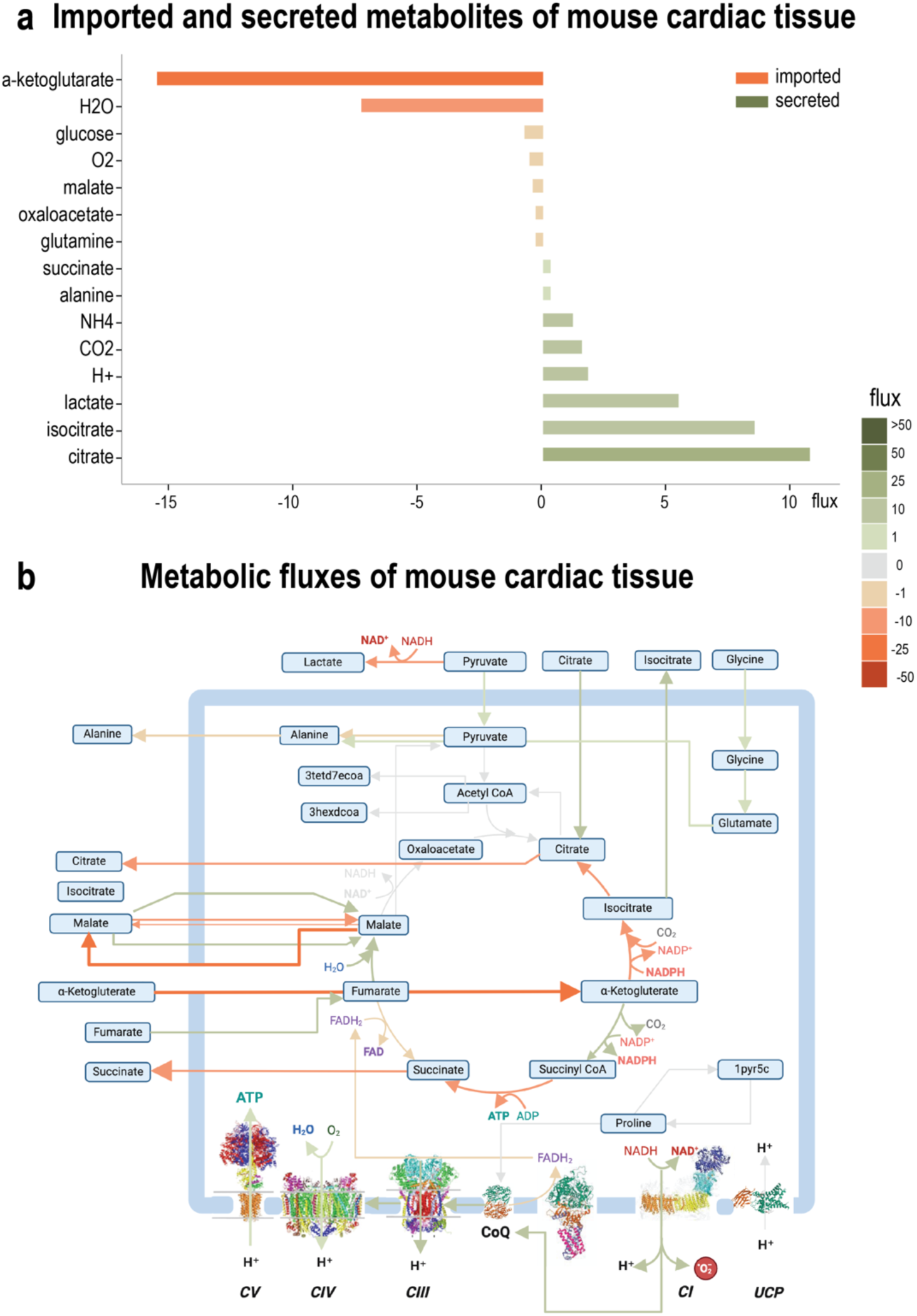
Metabolic flux prediction from mitoMammal following integration of mouse proteomic data isolated from cardiac tissue and optimising ATP production. **(a)** Imported (orange bars) and secreted (green bars) metabolites were predicted to result in steady-state mitochondrial fluxes of mitochondria isolated from mouse cardiac cells following the integration of proteomic data. Imported metabolites by convention, are associated with negative fluxes whilst secreted metabolites are described with positive fluxes. **(b)** The predicted flux distribution describes an import of glycine into the mitochondria, along with the import of citrate, fumarate and ɑKG. *De novo* fatty acid synthesis occurs as a result of pyruvate conversion into acetyl-CoA.

Flux predictions (Figure 3b) revealed that the flux of pyruvate emerging from glycolysis was partitioned between lactate production in the cytoplasm, and pyruvate import into the mitochondria. This is in agreement with the literature that reports a mitochondrial involvement of lactate production (Flick and Konieczny, 2002). It is now understood that the shuttle of lactate from and between cardiomyocytes to other cells facilitates lactate supply to cells in need of lactate, and acquired lactate plays a plethora of important roles such as cell signalling (Daw *et al*., 2020), the regulation of cell proliferation (Liu *et al*., 2023) and development of organs and in the coordination of vascular development and progenitor cell behaviour in the developing mouse neocortex (Dong *et al*., 2022). TCA cycle fluxes were sustained by the import of citrate, ɑKG, fumarate and malate. Glycine was imported into the mitochondria and converted to glutamate. Glycine has been shown to protect against heart toxicity in mice (Shosha *et al*., 2023) which validates this prediction, and highlights the important role of glycine metabolism in cardiomyocytes (Quintanilla-Villanueva *et al*., 2024) in sustaining steady-state metabolism.

### Modelling mouse BA metabolism by integrating mouse proteomic data with mitoMammal

We next wanted to show the predictive power and usability of mitoMammal to predict mouse mitochondrial metabolism in a BA cell by integrating mitochondrial proteomic data extracted from a BA cell (Hansen *et al*., 2024). Following data integration, we then optimised flux towards the UCP1 reaction. From the model’s complement of 560 reactions, our modified E-Flux2 algorithm constrained 329 reactions. In order of decreasing flux magnitude, mitoMammal predicted the import of glucose, citrate, ɑKG, fumarate, O2, cysteine, acetoacetate, butanoic acid, hydrogen, aspartate, alanine and finally malate. Secreted metabolites consisted of CO2, succinate, isocitrate, NH4, thiosulfate, hexadecanoic acid and propanoate (Figure 4a).

**Figure 4:**
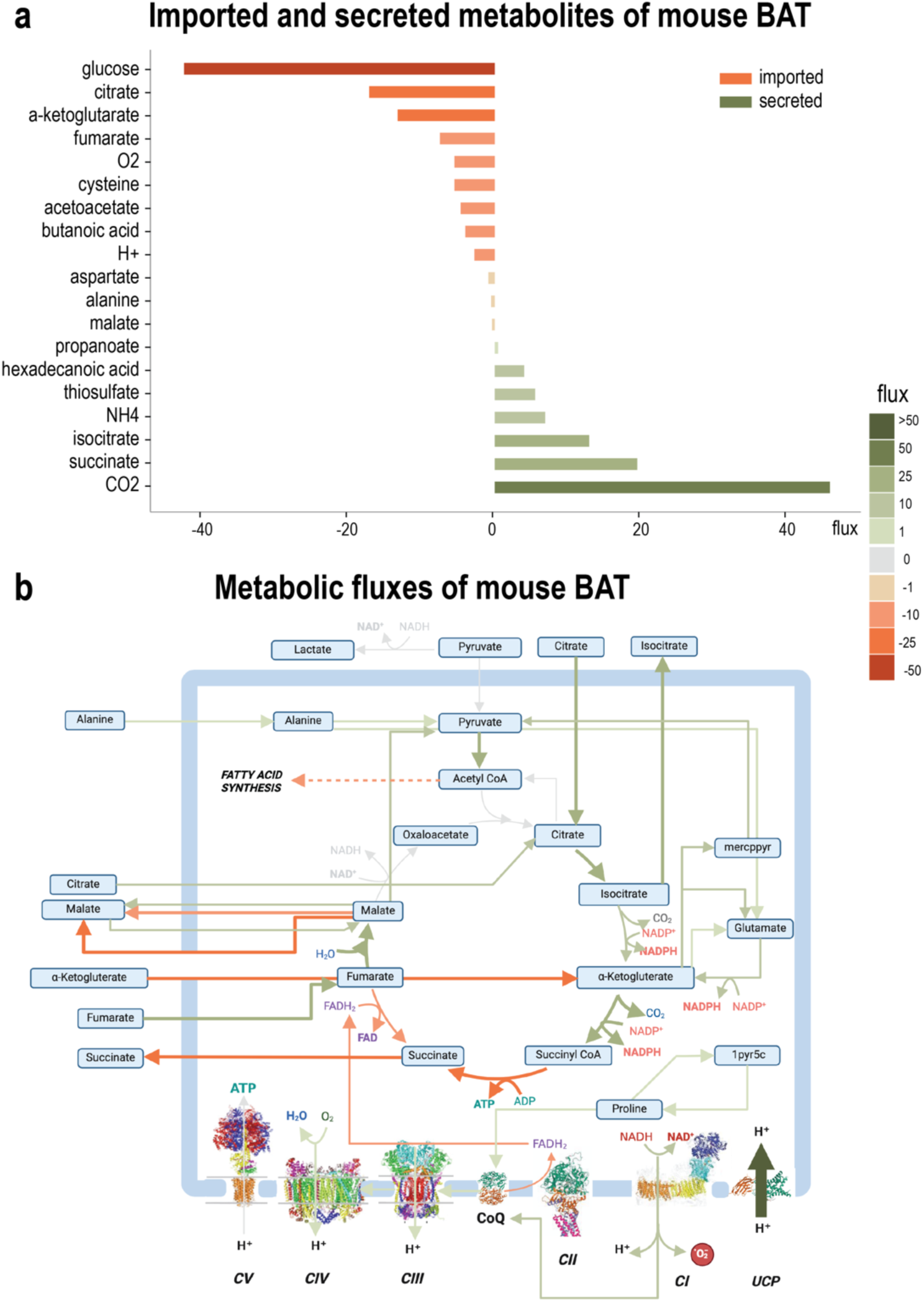
Proteomic data of mitochondria isolated from murine BAT was used to constrain mitoMammal. **(a)** Imported (orange bars) and secreted (green bars) metabolites were predicted to result in steady-state mitochondrial fluxes of mitochondria isolated from mice BAT cells following the integration of proteomic data. Imported metabolites by convention, are associated with negative fluxes whilst secreted metabolites are described with positive fluxes. **(b)** As a result of optimising the UCP1 reaction, the ETC is disengaged from ATP production emerging from OXPHOS. Steady-state TCA cycle fluxes are established by the import of alanine, citrate, fumarate, and ɑKG. *De novo* fatty acid synthesis occurs as a result of pyruvate conversion into acetyl-CoA. mitoMammal also predicts the reduction of CoQ by proline, which is produced by 1-Pyrroline-5-Carboxylate (1pyr5c).

In this simulation, citrate, fumarate, ɑKG and to a lesser extent, malate were predicted to be imported into the mitochondria to establish steady-state TCA cycle fluxes. Imported ɑKG was metabolised into succinyl-CoA within the TCA, and into 3-Mercaptopyruvic acid (mercppyr) exterior of the TCA cycle which then fed into pyruvate metabolism. Pyruvate metabolism was also established by the import of alanine and with conversion of malate into pyruvate. The majority of pyruvate was converted into acetyl-CoA and with the addition of citrate, channelled flux towards fatty acid synthesis and the export of hexadecanoic acid. Citrate was imported into mitochondria and assimilated into the TCA cycle, and upon conversion to Isocitrate, was then partially exported from mitochondria.

Complex I (CI) was predicted to be reduced by NADH emerging from the TCA cycle, which injected electrons into the ETC and reduced the CoQ complex. MitoMammal also predicted the reduction of CoQ with proline via the proline dehydrogenase reaction (PROD2mB, encoded by the PRODH gene). CII was predicted to operate in reverse and reduce fumarate leading to succinate production and its subsequent export. From CoQ, electrons were passed along the ETC towards CIII and CIV which produced PMF, however ATP synthase (CV) in this situation was predicted to be inactive, and the UCP1 reaction was active and carried the largest flux in this simulation. Mitochondrial uncoupling via UCP1 is a process that expends energy by oxidising nutrients to produce heat, instead of ATP. To better understand the role played by the UCP1 reaction in BAT tissue, we next examined the reactions that would consume the newly uncoupled protons after their re-entry into the mitochondria to identify novel functionalities of the UCP1 reaction in BAT. 20 proton-consuming reactions were identified and are shown in Supplementary Table S2.

The largest subset of these reactions performed metabolism of fatty acid and consisted of MECR14C and MECR16C which are responsible for fatty acid elongation of 3-Hydroxy Tetradecenoyl-7 Coenzyme A and 3-Hydroxyhexadecanoyl Coenzyme A respectively. Also belonging to this group were the reactions MTPC14, MTPC16, r0722, r0726, r0730, r0733 and r0791 all performed fatty acid oxidation roles and released NADH within the mitochondria. The remaining reactions of this subset (r0633, r0638, r0735) all consume mitochondrial NADP. 4 more reactions performed metabolite transport functions with a citrate-carrying reaction (r0917b) carrying the greatest flux of this analysis. This reaction exports isocitrate and protons in exchange for citrate import. The model predicted flux associated with the characterised mitochondrial carrier responsible for the export of phosphate and photons (Plt2mB) out of the mitochondria. The citrate-malate antiporter (CITtamB) was also predicted to be active in exporting malate and protons in exchange for the import of citrate. Uncoupled protons were also predicted to leak out of the mitochondria, described by the Hmt reaction.

A further subset of 3 reactions were implicated with amino acid metabolism. Within this subset, the reaction to carry the largest flux was 3-Mercaptopyruvate:Cyanide Sulfurtransferase (r0595m) which converts mercaptopyruvate and sulfate into pyruvate and thiosulfate. Also within this subset is the P5CRxm that involves the production of proline, and finally the methylmalonyl Coenzyme A decarboxylase reaction (MMCDm) which converted methylmalonyl-CoA into propionyl-CoA. The remaining reaction predicted to metabolise uncoupled protons was the CI reaction of the OXPHOS subsystem.

### Modelling brown adipocyte metabolism in humans

Next, we wanted to demonstrate mitoMammals ability of modelling human mitochondrial metabolism. To this end, we integrated transcriptomic data (Rao *et al*., 2023) from a BA that was differentiated from an IPSC and optimised the UCP1 reaction. This resulted in constraining 488 reactions out of the complement of 560 reactions. Analysis of the resulting fluxes revealed that 15 metabolites were predicted to be imported into mitoMammal to support steady-state mitochondrial BA metabolism. Similar to the mouse model, glucose, citrate, ɑKG, fumarate, O2, butanoic acid, H+, and malate were imported, however with different magnitudes. The largest flux was associated with H+ and not glucose. In addition, Glutamine, glycine, oxaloacetate, formate, Fe2, and argininosuccinate were imported. Similar excreted metabolites included CO2, succinate, NH4, hexadecanoic acid and propanoate. Opposed to the mouse model, alanine was exported, and not imported. In addition, the human model secreted H2O, proline, serine, malate, lactate and urea (Figure 5a).

**Figure 5:**
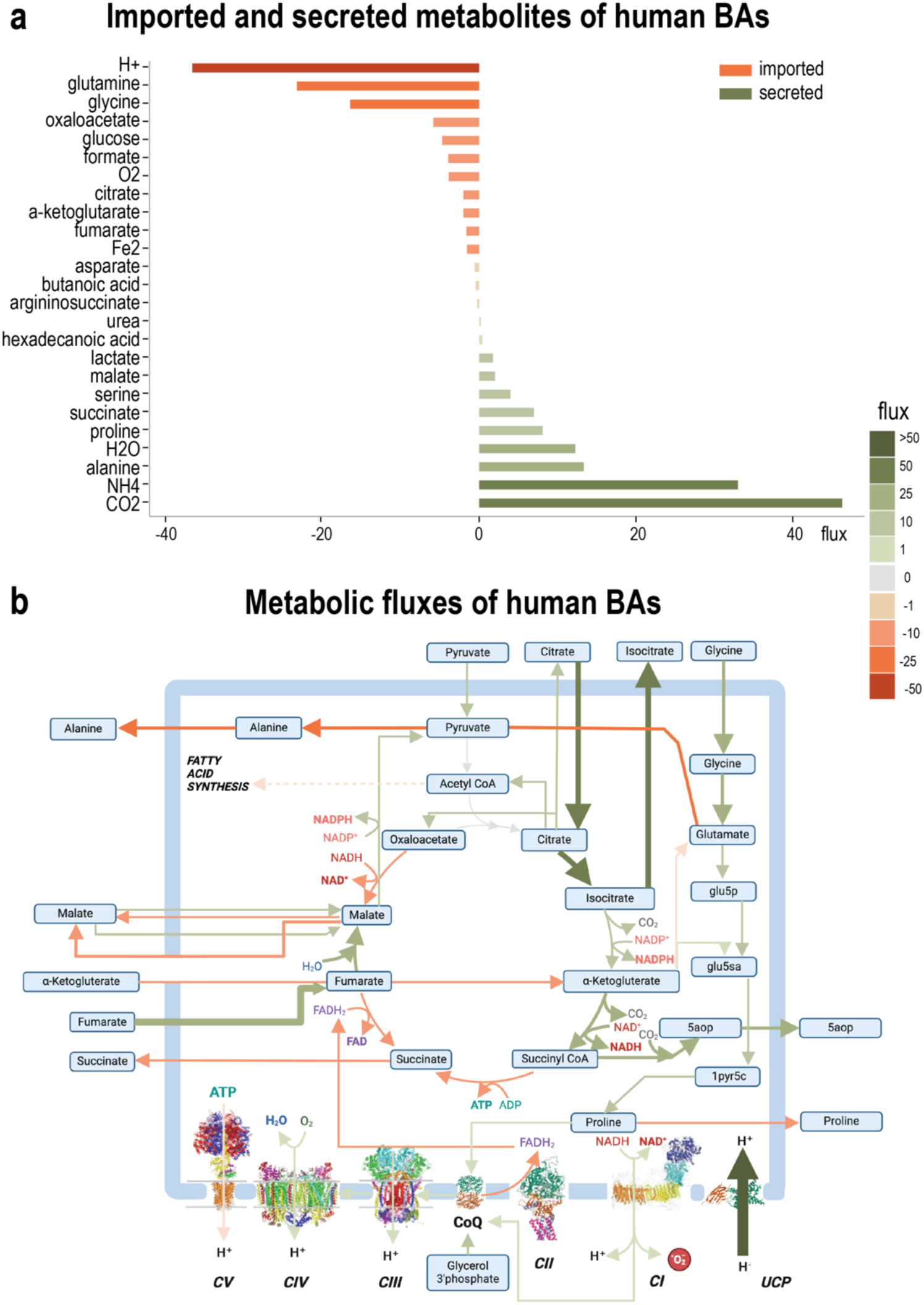
Flux distribution following the integration of transcriptomic data from a human brown adipocyte into mitoMammal. **(a)** Imported (orange bars) and secreted (green bars) metabolites predicted from steady-state mitochondrial fluxes of mitochondria from a human brown adipocyte following the integration of transcriptomic data. Imported metabolites by convention, are associated with negative fluxes whilst secreted metabolites are described with positive fluxes. **(b)** Metabolic fluxes are predicted to activate the UCP1 reaction which has the effect of uncoupling the ETC from OXPHOS to support steady-state metabolism during thermogenesis. Import of glycine, fumarate and ɑKG were predicted to sustain flux through the TCA cycle. Acetyl-CoA emerging from citrate import was then predicted to feed into *de novo* fatty acid synthesis. Metabolites exported out of the mitochondria were 5-aminolevulinic acid (5aop), proline, succinate, malate and alanine. Fluxes are highlighted in arrow thickness and colour; green colours: positive fluxes; orange colours: negative fluxes.

Similar to the mouse simulation, the import of citrate, fumarate, ɑKG and malate were predicted to contribute to sustaining steady-state TCA cycle fluxes. In human BAs, pyruvate emerging from glycolysis was predicted to be imported into the mitochondria and converted to alanine which was then, opposite to mouse BAs, exported out of the mitochondria. Citrate played a dual role and was also metabolised into acetyl-CoA which subsequently fed into endogenous fatty acid synthesis via acetyl-CoA, which agrees with the literature that describes mammalian BAT as possessing high endogenous fatty acid synthesis activity (Calderon-Dominguez *et al*., 2016; Schlein *et al*., 2021)(Figure 5b). In particular, the model predicted the synthesis and export of hexadecanoic acid and 5-Aminolevulinate (5aopm) from the mitochondria.

Fluxes through ETC were similar to Mouse BAT, except CV which this time was predicted to operate in reverse and consumed ATP. Similar to before, CII was predicted to operate in reverse. The model furthermore predicted the reduction of CoQ with proline via the proline dehydrogenase reaction (PROD2mB, encoded by the PRODH gene). PRODH forms part of the proline cycle that regenerates proline via pyrroline-5-carboxylate which, in contrast to the mouse model, leads to the subsequent export of proline. In addition, we predicted the reduction of CoQ by G3PDH, which was not predicted in the mouse simulation. The reaction carrying the greatest flux in this simulation was again the UCP1 reaction which uncoupled the ETC from ATP production. We then analysed all proton consumption reactions predicted to be active as a consequence of optimal UCP1 activity. All reactions that carry a flux greater than 0.01 are also shown in Supplementary Table S2.

As with the previous simulation of mouse BAT, the reaction to carry the greatest flux was attributed to the citrate-carrying reaction (r0917b). The largest subset of reactions was also implicated with the same fatty acid metabolic reactions as reported in the previous simulation, however carrying much reduced predicted fluxes. The next subset of reactions again, all involved transport functions with the first that exported phosphate and protons (Plt2mB) out of the mitochondria, and the citrate-malate antiporter (CITtamB). Uncoupled protons were also predicted to leak out of the mitochondria, as facilitated by the Hmt reaction. The final reaction predicted to be active in this simulation, which also was predicted to be active in the previous simulation, was the Pyrroline-5-Carboxylate Reductase reaction (P5CRxm).

The model also predicted a subset of reactions implicated with amino acid metabolism to be active in this simulation that was not predicted to be active in the mouse BAT simulation. Instead of predicting flux through r0595 and MMCDm reactions, the model instead predicted the consumption of uncoupled protons by 5-Aminolevulinate Synthase (ALASm) which metabolises glycine into 5aop_m. The final reaction of this subsystem involved the Glycine-Cleavage Complex (GCCAm) which converts glycine and lipoyl protein (lpro_m) into amino-methyl dihydrolipoyl protein (Alpro_m). The remaining reactions that were predicted to metabolise uncoupled protons following UCP1 optimisation and specific to the human simulations were Malate dehydrogenase (MDMm), a reaction belonging to folate metabolism (MTHFCm) and finally a reaction involved in the urea cycle (G5SDym).

## Discussion

We present mitoMammal, the first mitochondrial metabolic network reconstruction that serves for modelling mitochondrial metabolism for two species, mouse and human. MitoMammal contains two sets of GPR rules, one set of mouse genes, and another set of human genes, meaning the model can be constrained by integrating -omics data from these two organisms. Given the high similarity between mouse and human mitochondrial metabolism, we had the choice between two possible ways to model murine metabolism with -omics data-based constraints: either we would transform mouse gene identifiers to human and use the human mitoCore model 26/07*/2024 13*:09:00 for subsequent constraint-based modelling; or we could generate a mitochondrial metabolic model based on mitoCore that could be used for both species. We chose the latter, as it first makes the workflow for modelling mito-metabolism for the user straightforward; and second, it also allows the researcher to consider metabolic differences between the two organisms as each organism comes with its own set of GPR rules. We further added the DHODH reduction of CoQ following pyrimidine synthesis as this pathway was absent in mitoCore. As such, mitoMammal is the most comprehensive metabolic model of mammalian mitochondria to date.

To demonstrate the model’s ability to model mouse and human mitochondrial metabolism we first verified mitoMammals ability to capture realistic rates of ATP production. We then constrained mitoMammal by integrating proteomic data extracted from mouse cardiac tissue and optimised ATP production. Predicted fluxes included lactate production from pyruvate, the assimilation of pyruvate into the TCA cycle, the import of glycine into the mitochondria and the involvement of CV within OXPHOS to produce optimal ATP to support cardiomyocyte mitochondrial function. The model also predicted the reduction of CoQ by CI, yet fatty acid oxidation to support ATP synthesis was not predicted. These predictions are in agreement with data reported on immature cardiomyocytes, which express low levels of fatty acids and high levels of lactate in the blood that activates anaerobic glycolysis as the major source of ATP production (Karbassi *et al*., 2020).

Lactate is reported to fulfil important purposes that include providing an energy source for mitochondrial respiration, and being a major gluconeogenic precursor. As such, it is heavily involved in cellular signalling (Brooks, 2020). Several basic and clinical studies have revealed the role that lactate plays in heart failure with the consensus that high blood lactate levels indicate poor prognosis for heart failure patients (Zymliński *et al*., 2018). Current research on this topic aims to target lactate production, regulate lactate transport, and modulate circulating lactate levels in an attempt to find novel strategies for the treatment of cardiovascular diseases. The in-depth knowledge gained by metabolic modelling with mitoMammal could facilitate advances in this field.

To further demonstrate the usability of mitoMammal with alternative objective functions, and to highlight the ability of mitoMammal to model mouse and human metabolism, we integrated proteomic data extracted from the isolated mitochondria of mouse BAT (Figure 4) and integrated transcriptomic data of human BAs (Figure 5). For both simulations, we optimised the UCP1 reaction considering its central role in uncoupling electrons from the ETC. This leads to the dissipation of the PMF across the inner mitochondrial membrane which is essential for BAT function. Despite modelling two species with different -omics data sets, modelling BA metabolism with either human transcriptome or mouse proteome data resulted in several similar flux predictions. One such prediction relates to the metabolism of hexadecanoic acid, also known as palmitic acid, which has been shown to increase BA differentiation, decrease inflammation and improve whole-body glucose tolerance in mice (Unno *et al*., 2018) and humans (Wade *et al*., 2021). These data validate the predictions of hexadecanoic acid metabolism in both simulations.

Elevated levels of proline have been measured in mammalian BA tissue (Okamatsu-Ogura *et al*., 2020) and elevated levels of proline dehydrogenase have also been associated with BA differentiation, and thermogenesis and are correlated with UCP1 activity (Li *et al*., 2023). In both these simulations, mitoMammal indeed predicted proline reduction of CoQ via proline dehydrogenase, which is in line with these published data. Furthermore, it has been proposed that CoQ reduction by proline dehydrogenase activates ROS production which then activates signalling pathways that facilitate hormone-independent lipid catabolism and support adipose tissue thermogenesis (Lettieri Barbato and Aquilano, 2016; Chouchani *et al*., 2017).

All simulations predicted the reverse activity of CII. It has been experimentally demonstrated that CII can work in reverse in bacterial mitochondria (Maklashina *et al*., 1998) and mammalian mitochondria (Spinelli *et al*., 2021; Kumar *et al*., 2022). As reported by (Spinelli *et al*., 2021), fumarate was observed to be the final electron acceptor under oxygen-limiting situations. The work from (Kumar *et al*., 2022) further shows that CII is poised to operate either in the direction of succinate oxidation (forward activity) or fumarate reduction (reverse activity) at high hydrogen sulfate concentrations. They report that fumarate reduction results as a consequence of fumarate import into the TCA cycle which supports complex II reversal and leads to succinate accumulation. The predictions made by mitoMammal of fumarate import, its subsequent reduction as a consequence of CII reversal and succinate production and export agree with the reported literature. Succinate export from mitochondria from mice BAT has also been experimentally quantified using isotope tracing (Park *et al*., 2023) which further supports this novel prediction of the reverse activity of CII in supporting mice and human BAT metabolism.

We also observed differences in metabolic fluxes when comparing the predictions following human transcriptome integration and mouse proteomic integration. Following human transcriptomic integration, the model predicted the import of pyruvate into the mitochondria which was not predicted following mouse proteomic data integration. Instead, pyruvate was predicted to be converted to mercaptopyruvate (mercppyr). The simulation involving integrating human transcriptomic data also predicted the export of 5aopm which was not predicted when integrating mouse proteomic data. 5aopm is a precursor metabolite of the heme biosynthesis pathway and is required for adipocyte differentiation (Moreno-Navarrete and Fernández-Real, 2024). Disrupted heme biosynthesis in human and mouse adipocytes has been shown to result in decreased adipogenesis, impaired glucose uptake, and reduced mitochondrial respiration (Handschin *et al*., 2005; Moreno-Navarrete and Fernández-Real, 2024). These experimental discoveries of 5aopm therefore serve to further validate flux predictions following transcriptomic data integration and account for the misprediction associated with proteomic data integration. Alanine was also predicted to be imported into the mitochondria for the human transcriptome simulation, yet the mouse proteomic simulation predicted the export of alanine. Alanine import (Rodríguez-Martín and Remesar, 1991) and export (Frayn *et al*., 1991) into mammalian BAT tissue has been previously reported; however, the more comprehensive analysis reported by (Park *et al*., 2023) describes that alanine is an abundant circulating amino acid and functions as a nitrogen carrier where it is transported to the liver for nitrogen release. In their paper, the authors observe a net zero exchange flux and account for this to an equivalent uptake and release flux of alanine. As such, the model’s prediction of alanine import could be correct concerning mice metabolism (Figure 4a,b). Regarding human BAT metabolism, it is understood that accumulation of glutamate may increase the transamination of pyruvate to alanine (Borkum, 2023; Legendre *et al*., 2020), which mitoMammal predicts, but much less is known of the fate of alanine and further research is necessary to validate the specific prediction of the directionality of alanine metabolism in human BATs.

One remaining difference between the predictions is the activity of ATP synthase (CV) which was reported to operate in reverse following integration of RNA sequencing data and predicted to be inactive following integration of proteomic data from mice. MitoMammal represents the activity of CV as a Boolean representation of 14 genes that share an ‘AND’ relationship and so all 14 genes, or proteins need to be expressed to correctly produce all the individual subunits for a fully-functional enzyme. For these GPRs, all 14 RNA sequencing transcripts were quantified, and because of the known reversibility of CV, our adapted E-Flux2 algorithm constrained the upper and lower bounds that correlated to the lowest transcript level of these 14 genes. As a consequence, mitoMammal in this simulation predicted the reverse activity of CV. For the proteomic simulation however, two of the 14 proteins were not identified (ENSMUSG00000000563; ENSMUSG00000064357) and an additional 4 proteins (ENSMUSG00000006057; ENSMUSG00000062683; ENSMUSG00000018770; ENSMUSG00000018770) were recorded as zero counts. As such, CV in this simulation was effectively constrained to zero and took no part in sustaining metabolic flows. ATP synthase (CV) is well known to operate in reverse during a wide range of different physiological environments to generate a mitochondrial membrane potential through ATP hydrolysis (Acin-Perez *et al*., 2023; Junge and Nelson, 2015) and the capacity of ATP hydrolysis has been observed in mitochondria isolated from BAT from mice (Acin-Perez *et al*., 2023) and from humans (Harb *et al*., 2023). As such, these findings serve to validate the predictions made following the integration of human transcriptomic data, and highlight limitations of proteomic data in terms of missing data, as discussed in (Vanderaa and Gatto, 2021) and (Boys *et al*., 2023). We have indeed quantified this by observing the fact that integrating transcriptomic data resulted in constraining more reactions than proteomic data (489 reactions (BA, Human) vs. 329 (BAT mouse) or 330 (Cardiac mouse)).

Regarding the other reactions of the ETC, human BAT transcriptomic data integration predicted the reduction of CoQ by G3PDH and proline, yet for the mouse proteomic integrated simulation, proline reduced CoQ, and G3PDH was predicted to be inactive. G3PDH reduction of CoQ has been experimentally determined for BAT in both humans and mice (Banerjee *et al*., 2022; Oh *et al*., 2024). G3PDH is involved in the glycerol 3-phosphate shuttle, which, similarly to the MAS, shuttles reducing power in the form of NADH from the cytoplasm into the mitochondria. G3PDH then oxidises the imported NADH into NAD+ and releases an electron which reduces CoQ. Both mouse and human BAT express high levels of G3PDH, and knockout of G3PDH in both species are associated with metabolic Type 2 diabetes mellitus and obesity (Armani and Caprio, 2023; Banerjee *et al*., 2022; Brown *et al*., 2002). Given this, we believe the prediction of an inactive G3PDH flux in mice associated with proteomic data integration to be a misprediction as the G3PDH protein abundance (ENSMUSG00000026827; 331) was identified in the dataset, and so the upper bound was constrained to a corresponding positive value and the lower bound was constrained to zero. We therefore attribute this error to the FBA methodology that optimises metabolic fluxes to a given objective function. In this simulation, we optimised the UCP1 reactions and so the model calculated an optimal metabolic flow to the objective that did not involve the reduction of CoQ by G3PDH. This once more highlights the challenge associated with FBA of defining the correct biological objective reaction to optimise.

We have demonstrated that mitoMammal can be used with different objective functions which is a crucial step in constraint-based metabolic modelling (Dikicioglu *et al*., 2015; Schnitzer *et al*., 2022). In our simulations of heart metabolism, as a consequence of optimising maximum ATP production, metabolic flux was predicted to avoid the UCP1 reaction. This prediction has also been experimentally validated in the work of (Hansen *et al*., 2024) who show that the UCP1 protein is inactive for cardiac tissue, yet active in BAT cells which highlights the metabolic flexibility of mitochondria in supporting tissue-specific function.

## Notes

https://gitlab.com/habermann_lab/mitomammal

### Competing Interest Statement

The authors have declared no competing interest.

## References

Acin-Perez, R., et al. (2023) Inhibition of ATP synthase reverse activity restores energy homeostasis in mitochondrial pathologies. The EMBO Journal, 42, e111699.

Armani, A. and Caprio, M. (2023) Mineralocorticoid Receptor and Aldosterone: From Hydro-saline Metabolism to Metabolic Diseases. In, Caprio, M. and Fernandes-Rosa, F.L. (eds), Hydro Saline Metabolism, Endocrinology. Springer International Publishing, Cham, pp. 431–471.

Banerjee, R., et al. (2022) The mitochondrial coenzyme Q junction and complex III: biochemistry and pathophysiology. The FEBS Journal, 289, 6936–6958.

Ben Guebila, M. and Thiele, I. (2021) Dynamic flux balance analysis of whole-body metabolism for type 1 diabetes. Nat Comput Sci, 1, 348–361.

Bi, X., et al. (2022) Construction of Multiscale Genome-Scale Metabolic Models: Frameworks and Challenges. Biomolecules, 12, 721.

Borkum, J.M. (2023) The Tricarboxylic Acid Cycle as a Central Regulator of the Rate of Aging: Implications for Metabolic Interventions. Advanced Biology, 7, 2300095.

Borst, P. (2020) The malate–aspartate shuttle (Borst cycle): How it started and developed into a major metabolic pathway. IUBMB Life, 72, 2241–2259.

Boys, E.L., et al. (2023) Clinical applications of mass spectrometry-based proteomics in cancer: Where are we? Proteomics, 23, 2200238.

Brooks, G.A. (2020) Lactate as a fulcrum of metabolism. Redox Biology, 35, 101454.

Brown, L.J. et al. (2002) Normal Thyroid Thermogenesis but Reduced Viability and Adiposity in Mice Lacking the Mitochondrial Glycerol Phosphate Dehydrogenase. Journal of Biological Chemistry, 277, 32892–32898.

Brunk, E., et al. (2018) Recon3D enables a three-dimensional view of gene variation in human metabolism. Nat Biotechnol, 36, 272–281.

Cabbia, A., et al. (2021) Simulating Metabolic Flexibility in Low Energy Expenditure Conditions Using Genome-Scale Metabolic Models. Metabolites, 11, 695.

Calderon-Dominguez, M., et al. (2016) Fatty acid metabolism and the basis of brown adipose tissue function. Adipocyte, 5, 98–118.

Calvo, S.E. and Mootha, V.K. (2010) The Mitochondrial Proteome and Human Disease. Annu. Rev. Genom. Hum. Genet., 11, 25–44.

Chouchani, E.T., et al. (2017) Mitochondrial reactive oxygen species and adipose tissue thermogenesis: Bridging physiology and mechanisms. Journal of Biological Chemistry, 292, 16810–16816.

Daw, C.C., et al. (2020) Lactate Elicits ER-Mitochondrial Mg2+ Dynamics to Integrate Cellular Metabolism. Cell, 183, 474–489.e17.

Diaz-Cuadros, M., et al. (2023) Metabolic regulation of species-specific developmental rates. Nature, 613, 550–557.

Dikicioglu, D., et al. (2015) Biomass composition: the “elephant in the room” of metabolic modelling. Metabolomics, 11, 1690–1701.

Dong, X., et al. (2022) Metabolic lactate production coordinates vasculature development and progenitor behavior in the developing mouse neocortex. Nat Neurosci, 25, 865–875.

Duarte, N.C., et al. (2007) Global reconstruction of the human metabolic network based on genomic and bibliomic data. Proc. Natl. Acad. Sci. U.S.A., 104, 1777–1782.

Ebrahim, A., et al. (2013) COBRApy: COnstraints-Based Reconstruction and Analysis for Python. BMC Syst Biol, 7, 74.

Flatmark, T. and Pedersen, J.I. (1975) Brown adipose tissue mitochondria. Biochimica et Biophysica Acta (BBA) - Reviews on Bioenergetics, 416, 53–103.

Flick, M.J. and Konieczny, S.F. (2002) Identification of putative mammalian d-lactate dehydrogenase enzymes. Biochemical and Biophysical Research Communications, 295, 910–916.

Forman, H.J., et al. (2010) Signaling Functions of Reactive Oxygen Species. Biochemistry, 49, 835–842.

Frayn, K.N., et al. (1991) Amino acid metabolism in human subcutaneous adipose tissue *in vivo*. Clinical Science, 80, 471–474.

Fritzemeier, C.J., et al. (2017) Erroneous energy-generating cycles in published genome scale metabolic networks: Identification and removal. PLoS Comput Biol, 13, e1005494.

Handschin, C., et al. (2005) Nutritional Regulation of Hepatic Heme Biosynthesis and Porphyria through PGC-1α. Cell, 122, 505–515.

Hansen, F.M., et al. (2024) Mitochondrial phosphoproteomes are functionally specialized across tissues. Life Sci. Alliance, 7, e202302147.

Harb, E., et al. (2023) Brown adipose tissue and regulation of human body weight. Diabetes Metabolism Res, 39, e3594.

Hatefi, Y. (1985) THE MITOCHONDRIAL ELECTRON TRANSPORT AND OXIDATIVE PHOSPHORYLATION SYSTEM. Annu. Rev. Biochem., 54, 1015–1069.

Heinken, A., et al. (2021) Metabolic modelling reveals broad changes in gut microbial metabolism in inflammatory bowel disease patients with dysbiosis. npj Syst Biol Appl, 7, 19.

Junge, W. and Nelson, N. (2015) ATP Synthase. Annu. Rev. Biochem., 84, 631–657.

Karbassi, E., et al. (2020) Cardiomyocyte maturation: advances in knowledge and implications for regenerative medicine. Nat Rev Cardiol, 17, 341–359.

Karp, P.D., et al. (2023) The EcoCyc Database (2023). EcoSal Plus, 11, eesp-0002-2023.

Kasprzyk, A. (2011) BioMart: driving a paradigm change in biological data management. Database, 2011, bar049–bar049.

Keating, S.M., et al. (2020) SBML Level 3: an extensible format for the exchange and reuse of biological models. Molecular Systems Biology, 16, e9110.

Khodaee, S., et al. (2020) iMM1865: A New Reconstruction of Mouse Genome-Scale Metabolic Model. Sci Rep, 10, 6177.

Kim, M.K., et al. (2016) E-Flux2 and SPOT: Validated Methods for Inferring Intracellular Metabolic Flux Distributions from Transcriptomic Data. PLoS ONE, 11, e0157101.

Kim, M.K. and Lun, D.S. (2014) Methods for integration of transcriptomic data in genome-scale metabolic models. Computational and Structural Biotechnology Journal, 11, 59–65.

Kumar, R., et al. (2022) A redox cycle with complex II prioritizes sulfide quinone oxidoreductase-dependent H2S oxidation. Journal of Biological Chemistry, 298, 101435.

Kuznetsov, A.V., et al. (2009) The cell-type specificity of mitochondrial dynamics. The International Journal of Biochemistry & Cell Biology, 41, 1928–1939.

Legendre, F., et al. (2020) Biochemical pathways to α-ketoglutarate, a multi-faceted metabolite. World J Microbiol Biotechnol, 36, 123.

Lenaz, G., et al. (2007) The role of Coenzyme Q in mitochondrial electron transport. Mitochondrion, 7, S8–S33.

Lettieri Barbato, D. and Aquilano, K. (2016) Feast and famine: Adipose tissue adaptations for healthy aging. Ageing Research Reviews, 28, 85–93.

Li, F., et al. (2023) Proline hydroxylase 2 (PHD2) promotes brown adipose thermogenesis by enhancing the hydroxylation of UCP1. Molecular Metabolism, 73, 101747.

Liesa, M., et al. (2009) Mitochondrial Dynamics in Mammalian Health and Disease. Physiological Reviews, 89, 799–845.

Lister, A.L., et al. (2007) Integration of constraints documented in SBML, SBO, and the SBML Manual facilitates validation of biological models. Journal of Integrative Bioinformatics, 4, 252–263.

Liu, W., et al. (2023) Lactate regulates cell cycle by remodelling the anaphase promoting complex. Nature, 616, 790–797.

Majd, H., et al. (2018) Pathogenic mutations of the human mitochondrial citrate carrier SLC25A1 lead to impaired citrate export required for lipid, dolichol, ubiquinone and sterol synthesis. Biochimica et Biophysica Acta (BBA) - Bioenergetics, 1859, 1–7.

Maklashina, E., et al. (1998) Anaerobic Expression of *Escherichia coli* Succinate Dehydrogenase: Functional Replacement of Fumarate Reductase in the Respiratory Chain during Anaerobic Growth. J Bacteriol, 180, 5989–5996.

Marchiano, F., et al. (2022) The mitoXplorer 2.0 update: integrating and interpreting mitochondrial expression dynamics within a cellular context. Nucleic Acids Research, 50, W490– W499.

Martin, F.J., et al. (2023) Ensembl 2023. Nucleic Acids Research, 51, D933–D941.

Molinié, T., et al. (2022) MDH2 produced OAA is a metabolic switch rewiring the fuelling of respiratory chain and TCA cycle. Biochimica et Biophysica Acta (BBA) - Bioenergetics, 1863, 148532.

Moreno-Navarrete, J.M. and Fernández-Real, J.M. (2024) Iron: The silent culprit in your adipose tissue. Obesity Reviews, 25, e13647.

Mráček, T., et al. (2013) The function and the role of the mitochondrial glycerol-3-phosphate dehydrogenase in mammalian tissues. Biochimica et Biophysica Acta (BBA) - Bioenergetics, 1827, 401–410.

Nicholls, D.G. (1979) Brown adipose tissue mitochondria. Biochimica et Biophysica Acta (BBA) - Reviews on Bioenergetics, 549, 1–29.

Nicholls, D.G. (2005) Mitochondria and calcium signaling. Cell Calcium, 38, 311–317.

Nicholls, D.G. (2021) Mitochondrial proton leaks and uncoupling proteins. Biochimica et Biophysica Acta (BBA) - Bioenergetics, 1862, 148428.

Oh, S., et al. (2024) Glycerol 3-phosphate dehydrogenases (1 and 2) in cancer and other diseases. Exp Mol Med, 56, 1066–1079.

Okamatsu-Ogura, Y., et al. (2020) UCP1-dependent and UCP1-independent metabolic changes induced by acute cold exposure in brown adipose tissue of mice. Metabolism, 113, 154396.

Ong, W.K., et al. (2020) Model-driven analysis of mutant fitness experiments improves genome-scale metabolic models of Zymomonas mobilis ZM4. PLoS Comput Biol, 16, e1008137.

Orth, J.D., et al. (2010) What is flux balance analysis? Nat Biotechnol, 28, 245–248.

Pallag, G., et al. (2022) Proline Oxidation Supports Mitochondrial ATP Production When Complex I Is Inhibited. IJMS, 23, 5111.

Park, G., et al. (2023) Quantitative analysis of metabolic fluxes in brown fat and skeletal muscle during thermogenesis. Nat Metab, 5, 1204–1220.

Porter, C., et al. (2016) Human and Mouse Brown Adipose Tissue Mitochondria Have Comparable UCP1 Function. Cell Metabolism, 24, 246–255.

Quintanilla-Villanueva, G.E., et al. (2024) The Role of Amino Acid Glycine on Cardiovascular Health and Its Beneficial Effects: A Narrative Review. JVD, 3, 201–211.

Rao, J., et al. (2023) Reconstructing human brown fat developmental trajectory in vitro. Developmental Cell, 58, 2359–2375.e8.

Rodríguez-Martín, A. and Remesar, X. (1991) L-alanine transport in isolated cells of interscapular brown adipose tissue in rat. Bioscience Reports, 11, 65–71.

Sahu, A., et al. (2021) Advances in flux balance analysis by integrating machine learning and mechanism-based models. Computational and Structural Biotechnology Journal, 19, 4626–4640.

Schlein, C., et al. (2021) Endogenous Fatty Acid Synthesis Drives Brown Adipose Tissue Involution. Cell Reports, 34, 108624.

Schnitzer, B., et al. (2022) The choice of the objective function in flux balance analysis is crucial for predicting replicative lifespans in yeast. PLoS ONE, 17, e0276112.

Shosha, M.I., et al. (2023) Glycine protects against doxorubicin-induced heart toxicity in mice. Amino Acids, 55, 679–693.

Sigurdsson, M.I., et al. (2010) A detailed genome-wide reconstruction of mouse metabolism based on human Recon 1. BMC Syst Biol, 4, 140.

Smith, A.C., et al. (2017) MitoCore: a curated constraint-based model for simulating human central metabolism. BMC Syst Biol, 11, 114.

Smith, A.C. and Robinson, A.J. (2011) A metabolic model of the mitochondrion and its use in modelling diseases of the tricarboxylic acid cycle. BMC Syst Biol, 5, 102.

Spinelli, J.B., et al. (2021) Fumarate is a terminal electron acceptor in the mammalian electron transport chain. Science, 374, 1227–1237.

Suen, D.-F., et al. (2008) Mitochondrial dynamics and apoptosis. Genes Dev., 22, 1577–1590.

Takeda, Y. et al. (2023) Mitochondrial Energy Metabolism in the Regulation of Thermogenic Brown Fats and Human Metabolic Diseases. IJMS, 24, 1352.

Tanner, J.J., et al. (2018) The Proline Cycle As a Potential Cancer Therapy Target. Biochemistry, 57, 3433–3444.

Thiele, I., et al. (2013) A community-driven global reconstruction of human metabolism. Nat Biotechnol, 31, 419–425.

Tomi-Andrino, C., et al. (2022) Metabolic modeling-based drug repurposing in Glioblastoma. Sci Rep, 12, 11189.

Unno, Y., et al. (2018) Palmitoyl lactic acid induces adipogenesis and a brown fat-like phenotype in 3T3-L1 preadipocytes. Biochimica et Biophysica Acta (BBA) - Molecular and Cell Biology of Lipids, 1863, 772–782.

Vanderaa, C. and Gatto, L. (2021) Replication of single-cell proteomics data reveals important computational challenges. Expert Review of Proteomics, 18, 835–843.

Wade, G., et al. (2021) Lipid Transport in Brown Adipocyte Thermogenesis. Front. Physiol., 12, 787535.

Wagner, A., et al. (2021) Metabolic modeling of single Th17 cells reveals regulators of autoimmunity. Cell, 184, 4168–4185.e21.

Wang, G., et al. (2019) Regulation of UCP1 and Mitochondrial Metabolism in Brown Adipose Tissue by Reversible Succinylation. Molecular Cell, 74, 844–857.e7.

Westerhoff, H.V. (2023) On paradoxes between optimal growth, metabolic control analysis, and flux balance analysis. Biosystems, 233, 104998.

Williams, E.G., et al. (2018) Quantifying and Localizing the Mitochondrial Proteome Across Five Tissues in A Mouse Population. Molecular & Cellular Proteomics, 17, 1766–1777.

Zhou, Yue et al. (2021) DHODH and cancer: promising prospects to be explored. Cancer & Metabolism, 9, 22.

Zymliński, R., et al. (2018) Increased blood lactate is prevalent and identifies poor prognosis in patients with acute heart failure without overt peripheral hypoperfusion. European J of Heart Fail, 20, 1011–1018.

