## Supplementary Materials for "MitoMAMMAL: a genome scale model of mammalian mitochondria predicts cardiac and BAT metabolism"

**Supplementary Figure S1:**

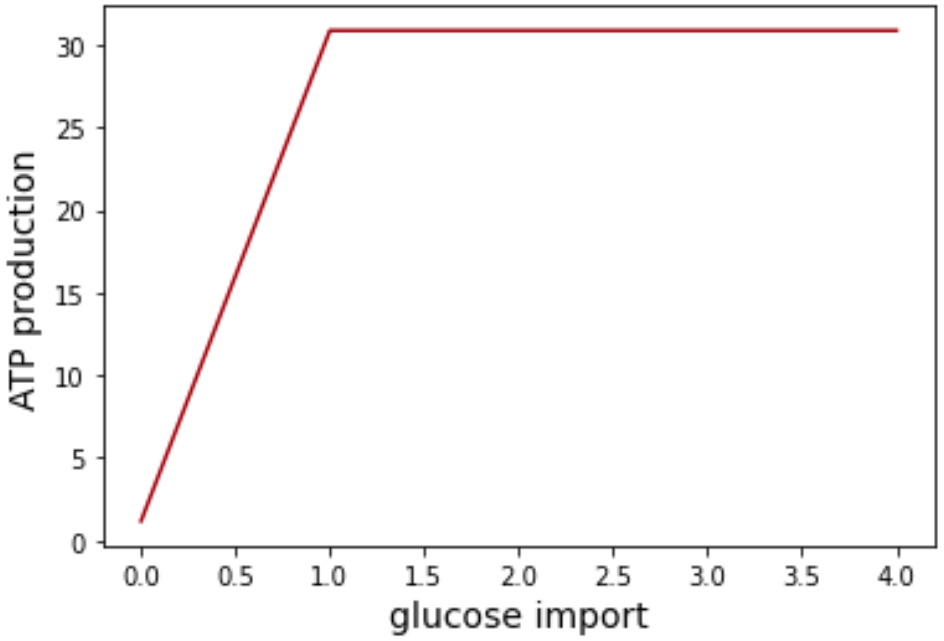


Supplementary Figure S1. 1 molecule of glucose import produces 31 molecules of ATP. By convention import fluxes are negative but to improve the comprehension we present the glucose import as positive values.

**Supplementary tables:**

**Supplementary Table S1.** A complete list of reactions, metabolites and associated fluxes presented in this manuscript are presented in Supplementary Tables S1a, S1b and S1c respectively (file Chapman_etal_SupplTableS1.xlsx).

**Supplementary Table S2:** reactions that consume hydrogen following the optimisation of the UCP1 reaction with integrated mouse proteomic data and human transcriptomic data.

| Reaction ID | Equation | Subsystem | Mouse BAT transcriptomic flux | Human BAT proteomic flux |
|---|---|---|---|---|
| ro917b | PMF_m + cit_c + h_m + icit_m <=> PMF_c + cit_m + h_c + icit_c | Transport | 12.88 | 25.67 |
| PIt2mB | 0.18 PMF_c + h_c + pi_c <=> 0.18 PMF_m + h_m + pi_m | Transport | 9.63 | 14.15 |
| Hmt | h_e <=> h_m | Transport | 6.80 | 0.43 |
| r0595 | h_m + mercppyr_m + so3_m --> pyr_m + tsul_m | Methionine and cysteine metabolism | 5.50 | 0.00 |
| CITtamB | PMF_m + cit_c + h_m + mal_L_m <=> PMF_c + cit_m + h_c + mal_L_c | Transport | 4.25 | 5.45 |
| MECR14C | 3tetd7ecoa_m + h_m + nadph_m --> nadp_m + tetd7ecoa_m | Fatty acid oxidation | 4.01 | 0.47 |
| MECR16C | 3hexdcoa_m + h_m + nadph_m --> nadp_m + pmtcoa_m | Fatty acid oxidation | 4.01 | 0.47 |
| MTPC14 | 3tetd7ecoa_m + coa_m + h2o_m + nad_m <=> accoa_m + ddcacoa_m + h_m + nadh_m | Fatty acid oxidation | 4.01 | 0.47 |
| MTPC16 | 3hexdcoa_m + coa_m + h2o_m + nad_m <=> accoa_m + h_m + nadh_m | Fatty acid oxidation | 4.01 | 0.47 |
| r0633 | nadp_m + occoa_m <=> HC01415_m + h_m + nadph_m | Fatty acid oxidation | 4.01 | 0.47 |
| r0638 | ddcacoa_m + nadp_m <=> dd2coa_m + h_m + nadph_m | Fatty acid oxidation | 4.01 | 0.47 |
| r0722 | HC01401_m + nad_m <=> 3oddcoa_m + h_m + nadh_m | Fatty acid oxidation | 4.01 | 0.47 |

| | | | | |
|---|---|---|---|---|
| r0726 | HC01403_m + nad_m <=> 3odcoa_m + h_m + nadh_m | Fatty acid oxidation | 4.01 | 0.47 |
| r0730 | HC01405_m + nad_m <=> HC01406_m + h_m + nadh_m | Fatty acid oxidation | 4.01 | 0.47 |
| r0733 | HC01407_m + nad_m <=> HC01408_m + h_m + nadh_m | Fatty acid oxidation | 4.01 | 0.47 |
| r0735 | dcacoa_m + nadp_m <=> dc2coa_m + h_m + nadph_m | Fatty acid oxidation | 4.01 | 0.47 |
| r0791 | HC01410_m + h_m + nadph_m --> HC01409_m + nadp_m | Fatty acid oxidation | 4.01 | 0.47 |
| CI | 3.996 PMF_m + h_m + nadh_m + 0.002 o2_m + 0.999 q10_m <=> 3.996 PMF_c + nad_m + 0.002 o2s_m + 0.999 q10h2_m | OXPHOS | 2.25 | 0.05 |
| P5CRxm | 1pyr5c_m + 2.0 h_m + nadh_m --> nad_m + pro_L_m | Arginine and proline metabolism | 1.76 | 18.11 |
| ALASm | gly_m + h_m + succoa_m --> 5aop_m + co2_m + coa_m | Glycine, serine, alanine, and threonine metabolism | 0 | 13.75 |
| G5SDym | glu5p_m + h_m + nadph_m <=> glu5sa_m + nadp_m + pi_m | Urea cycle | 0 | 8.74 |
| MTHFCm | h2o_m + methf_m <=> 10fthf_m + h_m | Folate metabolism | 0 | 4.37 |
| MDHm | mal_L_m + nad_m <=> h_m + nadh_m + oaa_m | TCA cycle | 0 | 2.1 |
| GCCam | gly_m + h_m + lpro_m <=> alpro_m + co2_m | Glycine, serine, alanine, and threonine metabolism | 0 | 0.07 |
